# Wnt7a-releasing synthetic hydrogel enhances local skeletal muscle regeneration and muscle stem cell engraftment

**DOI:** 10.1101/569921

**Authors:** Woojin M. Han, Mahir Mohiuddin, Shannon E. Anderson, Andrés J. García, Young C. Jang

**Author notes:** Co-corresponding authors: Andrés J. García, Ph.D. Executive Director, Parker H. Petit Institute for Bioengineering and Bioscience, Petit Director’s Chair in Bioengineering and Bioscience, Regents’ Professor, Woodruff School of Mechanical Engineering, Petit Institute for Bioengineering and Bioscience, Georgia Institute of Technology, 315 Ferst Drive NW, Atlanta, GA 30332-0363.; Young C. Jang, Ph.D. Assistant Professor, School of Biological Sciences, Petit Institute for Bioengineering and Bioscience, Georgia Institute of Technology, 315 Ferst Drive, 3304 Petit Biotechnology Building, Atlanta, GA 30332-0363.

## Abstract

Skeletal muscle possesses efficient ability to regenerate upon minor injuries, but its capacity to regenerate is severely compromised with traumatic injuries and muscle-associated diseases. Recent evidence suggests that skeletal muscle regeneration can be accelerated by transplantation of muscle satellite cells (MuSCs) or treatment with promyogenic factors, such as Wingless-type MMTV Integrated 7a (Wnt7a) protein. Although direct intramuscular injection is the simplest method to deliver MuSCs and Wnt7a for regenerative therapy, direct injection is not viable in many clinical cases where the structural integrity is severely compromised. To address this challenge, we engineered a synthetic poly(ethylene glycol) (PEG)-based hydrogel to facilitate the co-delivery of pro-myogenic factors, such as Wnt7a, and MuSCs to skeletal muscles affected by severe trauma or muscular dystrophies. Wnt7a release rate can be controlled by modulating the polymer density of the hydrogel, and this release rate can be further accelerated through the proteolytic degradation of the hydrogel. Treating cryo-injured tibialis anterior (TA) muscles with Wnt7a-loaded hydrogels resulted in an accelerated regenerative response, measured by increased muscle fiber cross-sectional area, bulk TA mass, and number of Pax7^+^ MuSCs at the injury site, compared to the TA muscles treated with Wnt7a-free hydrogels. Co-delivery of Wnt7a and primary MuSCs using the synthetic hydrogel to the cryo-injured TA muscles significantly increased cellular migration during the engraftment process. This work provides a synthetic biomaterial platform for advancing treatment strategies of skeletal muscle trauma and diseases, specifically in conditions where direct intramuscular injection may be challenging.

## INTRODUCTION

Skeletal muscle plays an indispensable role in locomotion and metabolism. Skeletal muscle regenerates efficiently following minor injuries, but its repair capacity is severely challenged with the onset of acute muscle trauma (e.g., laceration and volumetric muscle loss) and chronic muscle diseases (e.g., Duchenne muscular dystrophy) (1, 2). Inadequate muscle regeneration leads to incapacitating pathophysiologic consequences, such as aberrant inflammation, excessive fibrosis, and scar formation, ultimately resulting in permanent loss of muscle function, disability, and diminished quality of life (3, 4). Current muscle reconstruction strategies include surgical suturing and autologous muscle graft, but these approaches do not completely repair the injured muscle (3, 5). In addition, autologous muscle transplant causes donor site morbidity and is prone to graft failure, especially in aged patients with muscle wasting conditions (3). Effective therapeutic strategies for promoting skeletal muscle regeneration remain to be developed.

Recent studies have revealed that Wingless-type MMTV Integrated 7a (Wnt7a) protein, a ligand for Frizzled (Fzd) receptors, functions as a potent mediator of muscle growth and repair (6–10). Notably, Wnt7a induces myofiber hypertrophy through the Akt/mTOR protein synthesis pathway (8, 9), promotes symmetric expansion (6, 10), and migration of muscle satellite cells (MuSCs) through the planar cell polarity pathway involving Dishevelled 2 (Dvl2) and the small GTPase Rac1 (6, 7, 10). Furthermore, pre-treatment of MuSCs with Wnt7a significantly enhances MuSC dispersion and engraftment upon cellular transplantation (7). In a rodent model of Duchenne muscular dystrophy (DMD), administration of recombinant human Wnt7a into tibialis anterior muscles significantly promotes MuSC expansion, myofiber hypertrophy, and muscle strength, suggesting that Wnt7a serves as an effective pro-myogenic factor for stimulating muscle repair (9). Although Wnt7a treatment may also enhance local muscle regeneration upon injury, traumatic injuries are often characterized by a severe loss in tissue integrity (11, 12), rendering direct injection of Wnt7a and other therapeutics not feasible. Therefore, there is a need for a delivery vehicle that enables controlled administration of therapeutics to the injured or pathologic muscles where direct injection may not be applicable.

Hydrogel-based biomaterials offer a promising strategy to deliver therapeutics, including stem/progenitor cells and growth factors (13–15). These biomaterial systems enable precise control over the spatiotemporal release of therapeutic molecules and address clinical challenges associated with requiring high dosages, repeated administration, and short half-lives of protein-based agents (13). Recently, we engineered a synthetic bioactive hydrogel based on a four-arm poly(ethylene glycol) (PEG) macromer with its ends functionalized with maleimide groups (PEG-4MAL) for delivering MuSCs to dystrophic and aged skeletal muscles with comorbid trauma (16). The engineered PEG-4MAL hydrogel provides several advantages for treating muscle trauma and disorders, including its fully synthetic nature that increases its translational potential, and ability to systematically control the structure, physical and biochemical properties, and degradability (16–18). Furthermore, the engineered PEG-4MAL hydrogel efficiently adheres to biological tissue through maleimide-thiol interactions (Michael-type addition) at physiological conditions (18), making the system suitable for locally delivering cells and therapeutics to the injured or pathologic muscles where direct injection is impractical. We also demonstrated that the engineered PEG-4MAL hydrogel supports primary MuSC function, and MuSC delivery using the material significantly improves their engraftment efficiency compared to hydrogel-free and collagen gel controls (16). However, the effect of co-delivering pro-myogenic growth factors, such as Wnt7a, with MuSC using the engineered PEG-4MAL hydrogel on stimulating muscle regeneration has not yet been evaluated.

In this study, we engineered a synthetic PEG-4MAL hydrogel to facilitate the co-delivery of Wnt7a and MuSCs for accelerated skeletal muscle healing. We demonstrate that the release rate of Wnt7a can be controlled by modulating the hydrogel polymer density, and the hydrogel-released Wnt7a retains its bioactivity both *in vitro* and *in vivo*. Supramuscular delivery of Wnt7a-releasing hydrogel to the freeze-injured tibialis anterior (TA) muscle accelerates the regenerative response by promoting muscle fiber hypertrophy and increasing endogenous Pax7^+^ MuSCs at the site of injury. Finally, we demonstrate that the co-delivery of Wnt7a and MuSCs using the engineered PEG-4MAL hydrogel synergistically enhances the migration capacity of donor MuSCs into the injured TA muscle. Collectively, we establish codelivery of Wnt7a and MuSCs using the engineered synthetic PEG-4MAL hydrogel as a potential therapeutic strategy to locally treat skeletal muscle injuries and disorders where direct injection may not be applicable.

## MATERIALS AND METHODS

### Animals

All live animal procedures were conducted under the approval of the Institutional Animal Care and Use Committee of the Georgia Institute of Technology. Mice were housed, maintained, and bred in the Physiological Research Laboratory Animal Facility of the Georgia Institute of Technology. *C57Bl/6* and *mdx* (*B6Ros.Cg-DMD*^*mdx-^4^CV*^*/J*) mice were obtained from the Jackson Laboratory. CAG-luc-Ka-β-actin-EGFP mice were generated by crossing *C57Bl/6-*β*-actin-EGFP* (kindly provided by Amy Wagers, Harvard University) and *FVB-Tg(CAG-luc,-GFP)L2G85Chco/J* (acquired from the Jackson Laboratory) mice. These mice were back-crossed for more than 5 generations and maintained in *C57Bl/6* background. Both male and female mice were used in a randomized manner.

### Synthesis of Wnt7a-loaded hydrogel

Four-arm PEG-4MAL macromer (20 kDa, Laysan Bio) was reconstituted in 1X phosphate-buffered saline (PBS) containing 10 mM HEPES (pH 7.4). Cell adhesive peptides (GRGDSPC, or scrambled GRDGSPC; >95% purity; GenScript) were dissolved in PBS containing 10 mM HEPES. To generate peptide-functionalized PEG-4MAL precursors, cell adhesive peptides were mixed with the PEG-4MAL solution. Subsequently, recombinant human Wnt7a (R&D systems) reconstituted in PBS was added to the PEG-4MAL precursor solution. To synthesize Wnt7a-loaded hydrogel, protease-degradable cross-linking peptide (GCRD<u>VPM</u>SMRGGDRCG; >95% purity; GenScript) dissolved in PBS containing 10 mM HEPES was mixed with the PEG-4MAL precursor solution containing Wnt7a and polymerized at 37 °C for 5 min.

### Release assay

Wnt7a was labeled with AlexaFluor 488 NHS ester (ThermoFisher Scientific) per the manufacturer’s instructions. 1 µg labeled-Wnt7a was encapsulated in a 15 µL PEG-4MAL hydrogel and subsequently submerged in 1 mL PBS. Release kinetics of Wnt7a from hydrogels was fluorescently quantified over time at 37 °C by sampling the bathing solution in a 24-well microvolume microplate and SpectraMax M3 reader (Molecular Devices). For each time point, PBS (blank control) and free 1 µg labeled-Wnt7a in 1 mL PBS (100% control) were also assayed. To degrade Wnt7a-loaded hydrogels, the hydrogels were incubated in collagenase I (3.9 U/mL in PBS; Worthington Biochemical Corporation) at 37 °C.

### In vitro bioactivity assay

C2C12 myoblasts were cultured in 1.5 mL growth media (Dulbecco’s modified Eagle’s medium, DMEM containing 100 U/mL penicillin/streptomycin and 10% fetal bovine serum [FBS]) in 12-well plates until confluency. Subsequently, the growth media was replaced with 1.5 mL of differentiation media (DMEM containing 100 U/mL penicillin/streptomycin and 2% horse serum) containing Wnt7a (75 ng)-loaded PEG-4MAL hydrogels (4%, 6%, 8% w/v), free Wnt7a (75 ng), or PBS control. After 5 days of differentiation, myotubes were fixed using 4% paraformaldehyde for 20 min at room temperature, washed, treated with blocking/permeabilizing buffer (2% bovine serum albumin [BSA], 0.5% goat serum, and 0.5% Triton X-100 in PBS), and stained for myosin heavy chain (1:200; MF20, Developmental Studies Hybridoma Bank) overnight at 4 °C. The myotubes were washed 3 times using PBS containing Tween-20 (0.1% v/v) and incubated with AlexaFluor 488-conjugated goat anti-mouse IgG (1:250; ThermoFisher) for 1 hour at room temperature. Samples were washed 3 times using PBS containing Tween-20 (0.1% v/v) and imaged using an epifluorescent microscope (Zeiss Observer D1). To assess the degree of hypertrophy, myotube diameters were measured using Fiji ImageJ 1.52e (19).

### PEG-4MAL macromer muscle adhesion assay

Scrambled RDG peptides (GRDGSPC; >95% purity; GenScript) were labeled with DyLight 755 NHS ester (ThermoFisher Scientific) per the manufacturer’s instructions. DyLight 755-labeled RDG-functionalized PEG-4MAL precursors (remaining maleimide concentration = 6.27 × 10^−3^ mol/mL) were produced as described above. To generate maleimide-quenched precursors, excess L-cysteine (Millipore Sigma) dissolved in PBS was added to the PEG-4MAL solution. 15 µL of PBS, macromer solution with quenched maleimides, or macromer solution with reactive maleimides was pipetted onto the TA surfaces of freshly sacrificed mdx-4CV mice. Treated TA muscles were harvested and washed for 5 min twice in PBS. Fluorescence measurement of the TAs was acquired using the IVIS SpectrumCT *in vivo* imaging system (PerkinElmer).

### MuSC isolation

Primary MuSCs were isolated as described previously (16). Briefly, hindlimb muscles from 2-4-month-old mice were minced and digested in DMEM containing collagenase type II (0.2%; Worthington Biochemical Corporation) and dispase II (2.5 U/mL; ThermoFisher) for 90 min at 37 °C on a shaker. The muscle digest was triturated every 30 min using an FBS-coated serological pipette. Upon digestion, the muscle digest was diluted by adding a two-part volume of 20% FBS in F10 media. The resulting solution was filtered through a 70 µm cell strainer. Cell pellets were obtained by centrifuging (300*g*) the solution for 5 min at 4 °C. The cell pellets were retrieved by removing supernatant and further washed using 2% FBS in Hank’s balanced salt solution (HBSS) through 5 min centrifugation (300*g*) at 4 °C. To stain the cells with primary antibodies, a cocktail of primary antibodies (1:100) was added to the cell suspension solution in 2% FBS in HBSS and incubated for 30 min on ice. The primary antibody cocktail consists of phycoerythrin (PE)-conjugated anti-mouse CD29 (13.3%; BioLegend), allophycocyanin (APC)-conjugated anti-mouse Sca1 (13.3%; BioLegend), CD45 (13.3%; BioLegend), CD11b (13.3%; BioLegend), CD31 (13.3%; BioLegend), Ter119 (13.3%; BioLegend), and biotinylated anti-mouse CD184 (20.2%; BD Pharmingen). The cells were washed in 2% FBS in HBSS through 5 min centrifugation (300*g*) at 4 °C. The cells were further incubated with streptavidin PE-Cy7 (1:100; eBioscience) for 20 min on ice and washed as described above. Immediately before cell sorting, propidium iodide (1:1000; ThermoFisher) was added to the cell solution. Propidium iodide^-^, CD29^+^, CD184^+^, Sca1^-^, CD31^-^, CD45^-^, CD11b^-^, Ter119^-^ MuSCs were sorted via fluorescence-activated cell sorting (BD FACSAria III Cell Sorter).

### 3D MuSC culture, proliferation, and migration assay

Freshly isolated MuSCs were encapsulated in 4% 20 kDa PEG-4MAL hydrogels (1.0 mM RGD, 50 ng/mL Wnt7a, VPM crosslinked) as described above. MuSCs were cultured in growth media (F10 containing 100 U/mL penicillin/streptomycin, 1X GlutaMAX, and 20% horse serum). For EdU staining, cells were incubated in 10 µM EdU-containing growth media for 5 hours. Subsequently, the cells were fixed in 4% paraformaldehyde for 20 min at room temperature and washed three times using PBS. The cells were blocked and permeabilized using PBS containing 2% BSA, 0.5% goat serum, and 0.5% Triton X-100 for 1 hour at room temperature. EdU detection was conducted per the manufacturer’s instructions (ThermoFisher). Cell nuclei were stained with Hoechst (1:1000) prior to z-stack imaging using a confocal microscope (Nikon Eclipse Ti-E C2+). The number of nuclei and EdU^+^ nuclei were automatically counted using CellProfiler (20). Total colony area per volume was quantified using ImageJ. To assess MuSC migration, 4% 20 kDa PEG-4MAL hydrogels (1.0 mM RGD, VPM crosslinked) containing Wnt7a (50 ng) or PBS were cast in angiogenesis plate wells (ibidi), and freshly isolated 10,000 GFP^+^ MuSCs were seeded on top of the hydrogels. 1,360 µm-thick z-stack projections were taken, from the top of the hydrogel, using a confocal microscope (Nikon Eclipse Ti-E C2+). Cell migration distance was quantified using ImageJ.

### Freeze injury and Wnt7a/MuSC delivery

Mice were anesthetized by inhalation of 2.5% isoflurane. Sustained-release buprenorphine (0.8 mg/kg) was subcutaneously administered immediately after induction. The hindlimb hair was depilated to expose the skin. To expose the TA muscle, an incision was made through the skin and fascia between the TA and gastrocnemius muscles on the lateral side. A liquid nitrogen-cooled metal probe was positioned on the TA surface for 10 sec to induce freeze injury. Upon muscle recovery, 15 µL PEG-4MAL hydrogel (4% 20 kDa PEG-4MAL, 1.0 mM RGD, VPM crosslinked) containing 2.5 µg Wnt7a or PBS was cast on the injured TA of *C57Bl/6* mice. For hydrogel-mediated co-delivery of Wnt7a and freshly isolated GFP^+^ MuSCs, 15 µL PEG-4MAL hydrogel (3% 20 kDa PEG-4MAL, 1.0 mM RGD, VPM crosslinked) containing 300 ng Wnt7a with MuSCs or PBS with MuSCs was cast on the injured TA of *mdx* mice. The skin incision was closed using degradable surgical suture (Vicryl suture, Ethicon). All procedures were conducted using aseptic techniques.

### Tissue histology and immunostaining

Hindlimbs were fixed in 4% paraformaldehyde for 1 hour at room temperature. Fixed hindlimbs were washed in PBS three times and incubated in 20% sucrose (w/v in PBS) overnight at 4 °C. Dissected TA muscles were frozen in liquid nitrogen-cooled 2-methylbutane. 10 µm-thick tissue sections were acquired through cryo-sectioning. Prior to tissue staining, the sectioned tissues were treated with blocking/permeabilizing buffer (2% BSA, 0.5% goat serum, and 0.5% Triton X-100 in PBS) for 1 hour at room temperature.

To stain for Pax7, TA muscles were frozen unfixed. 10 µm-thick tissue sections were obtained and fixed using 4% paraformaldehyde for 8 min. Heat activated antigen retrieval was performed by incubating the tissues in citrate buffer (pH 6.0) and treating in a high-pressure cooker for 10 min. The tissue sections were washed using PBS, blocked using blocking/permeabilizing buffer (2% BSA, 0.5% goat serum, and 0.5% Triton X-100) for 1 hour at room temperature, and finally incubated in blocking/permeabilizing buffer containing AffiniPure Fab goat anti-mouse-IgG (1:25; Jackson ImmunoResearch Laboratories, Inc.) for 1-2 hours at room temperature

The following primary and secondary antibodies were used to stain tissue sections in this study: rabbit polyclonal anti-dystrophin (1:200; Abcam), Pax7 (1:80, Developmental Studies Hybridoma Bank), Alexa Fluor 488-conjugated anti-GFP (1:250; ThermoFisher), Alexa Fluor 555 goat anti-mouse IgG (1:250; ThermoFisher), Alexa Fluor 488 goat anti-rabbit IgG (1:250; ThermoFisher). Hoechst 33342 (1:1000; Thermo Fisher) was used to counterstain cell nuclei.

### Pax7^+^ MuSC and myofiber cross-sectional area quantification

To quantify Pax7^+^ MuSCs, tissue sections were imaged and manually quantified in a blinded manner, where the slide labels were physically masked using a nontransparent tape by a different researcher who did not partake in the imaging and quantification process. The number of Pax7^+^ MuSCs were scored from 8 images per muscle. Myofiber cross-sectional area was automatically quantified using ImageJ. Briefly, the dystrophin images were converted 8-bit and automatic threshold was applied. Myofiber cross-sectional areas were then automatically quantified using the Analyze Particle function.

### Statistical analyses

Statistical tests were conducted using GraphPad Prism 8. Normality was tested using the Shapiro-Wilk test. Two-way repeated measures ANOVA with Tukey’s multiple comparisons tests were performed on the Wnt7a release data. Two-way ANOVA with Sidak’s multiple comparisons tests was performed to myogenic colony formation and EdU incorporation assay data. For all other multiple group comparisons, one-way ANOVA with Tukey’s tests or Kruskal-Wallis test with Dunn’s tests were performed based on data normality. For the paired comparisons, two-tailed paired t-tests were conducted. Two-tailed unpaired t-tests were performed for all other two-group comparisons. Significance levels were set at \**p*<0.05, \*\**p*<0.01, \*\*\**p*<0.01, \*\*\*\**p*<0.001.

## RESULTS

### Synthetic PEG-4MAL hydrogel for controlled-release of Wnt7a

To engineer a Wnt7a-releasing hydrogel, we encapsulated Wnt7a in a hydrogel composed of a four-arm poly(ethylene glycol) (PEG) macromer with its ends functionalized with maleimide groups (PEG-4MAL). Previously, we demonstrated that PEG-4MAL hydrogels can be customized to effectively deliver growth factors for various tissue regeneration applications, including vascular endothelial growth factor for pancreatic islet vascularization (21–23) and bone morphogenetic protein-2 for bone regeneration (24). Furthermore, PEG-4MAL hydrogels exhibit excellent biocompatibility *in vivo* (16, 18, 23, 25). In skeletal muscles, the engineered PEG-4MAL platform supports primary MuSC function and serves as an effective delivery vehicle for MuSC transplantation (16). In this system, the maleimide functional groups enable stoichiometric incorporation of thiol-presenting cell-adhesive ligands through Michael-type addition (Fig. 1a). Functionalized macromers can then be polymerized using cysteine-flanked protease-degradable peptides, which results in a hydrogel with a highly defined network structure that exhibits “on-demand” degradability (Fig. 1a). Furthermore, hydrogel mesh size, which governs solute diffusion through the hydrogel network, can be decreased by increasing the polymer density, permitting precise control of growth factor release rate.

**Figure 1.**
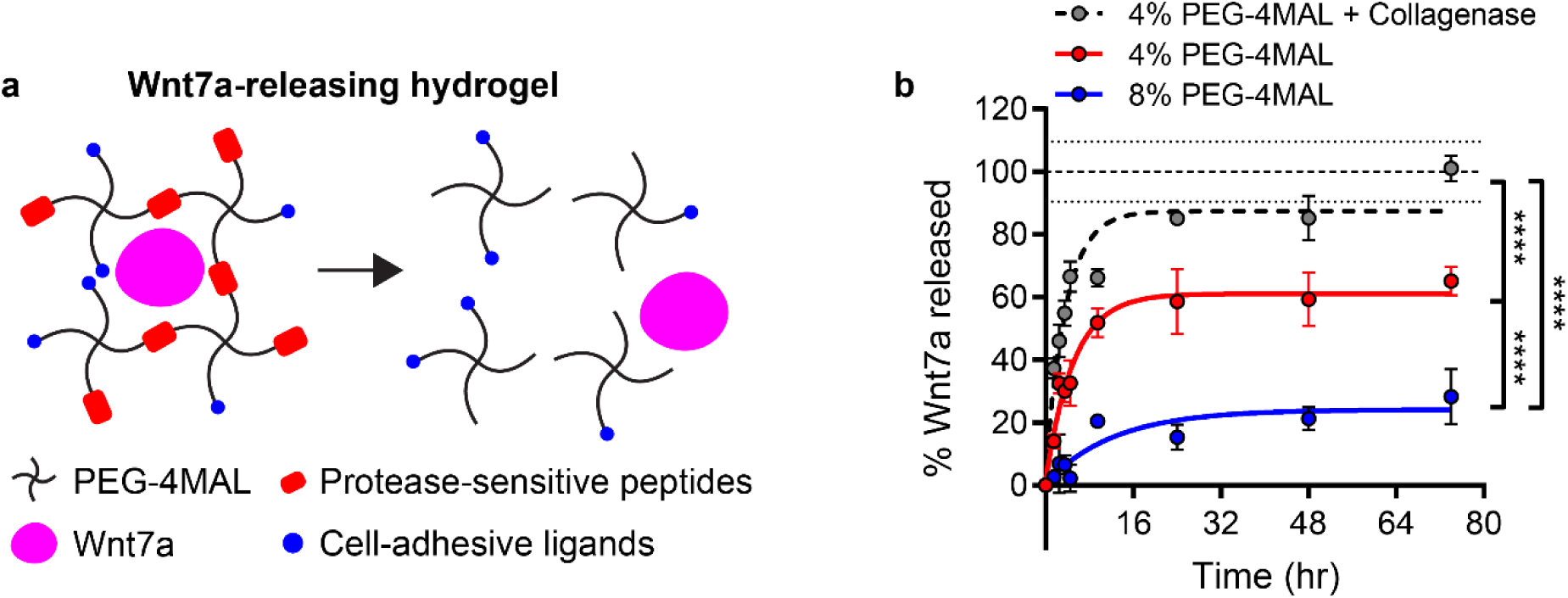
Synthetic PEG-4MAL hydrogel for controlled release of Wnt7a. **(a)** Schematic diagram of Wnt7a-releasing PEG-4MAL hydrogel. **(b)** Passive and proteolytic Wnt7a release profile with one-phase association fit. Extra sum of squares F-test for comparing curve fit. Mean ± SEM. n=4 hydrogels/condition. \*\*\*\**p*<0.0001.

Wnt7a release rate can be modulated by encapsulating in PEG-4MAL hydrogels with different polymer densities. The release rate of Wnt7a through passive diffusion is significantly delayed in 8% (w/v) PEG-MAL hydrogel compared to 6% PEG-4MAL hydrogel (Fig. 1b; *p*<0.0001) in the absence of collagenase; this result is expected based on the tighter network mesh for the higher polymer density gels. The release rate of Wnt7a is further increased in presence of protease (i.e., 3.9 U/mL of collagenase type I; Fig. 1b; *p*<0.0001), indicating that the release rate can be further enhanced through proteolytic degradation of the hydrogel.

### Hydrogel-released Wnt7a retains its bioactivity *in vitro*

To determine whether hydrogel-released Wnt7a retains its bioactivity, we assessed the capacity of hydrogel-released Wnt7a to promote *in vitro* myotube hypertrophy (Fig. 2a). In differentiated myotubes, Wnt7a binds to Fzd7 and activates the Akt/mTOR protein synthesis pathway to induce myotube hypertrophy (8). Significant hypertrophy of C2C12 myotubes was observed with treatment with free Wnt7a (Fig. 2b, c, g; *p*<0.05) compared to PBS control. Treating C2C12 myotubes with Wnt7a-releasing 4% PEG-4MAL hydrogel resulted in a comparable degree of hypertrophy as free Wnt7a (Fig. 2c, d, g; *p*>0.05). Treatment with Wnt7a-loaded 6% and 8% PEG-4MAL hydrogels resulted in decreased levels of myotube hypertrophy, compared to both free Wnt7a and 4% PEG-4MAL conditions (Fig. 2c, d, e, f, g; *p*<0.05). The distribution of myotube diameter increased with decreasing hydrogel polymer density (Fig. 2h), further suggesting that the effect of delivered Wnt7a can be temporally modulated by altering the Wnt7a release kinetics (Fig. 1b). Collectively, these data indicate that hydrogel-released Wnt7a retains its bioactivity *in vitro*, and further suggest that the release rate of Wnt7a modulates myotube hypertrophy.

**Figure 2.**
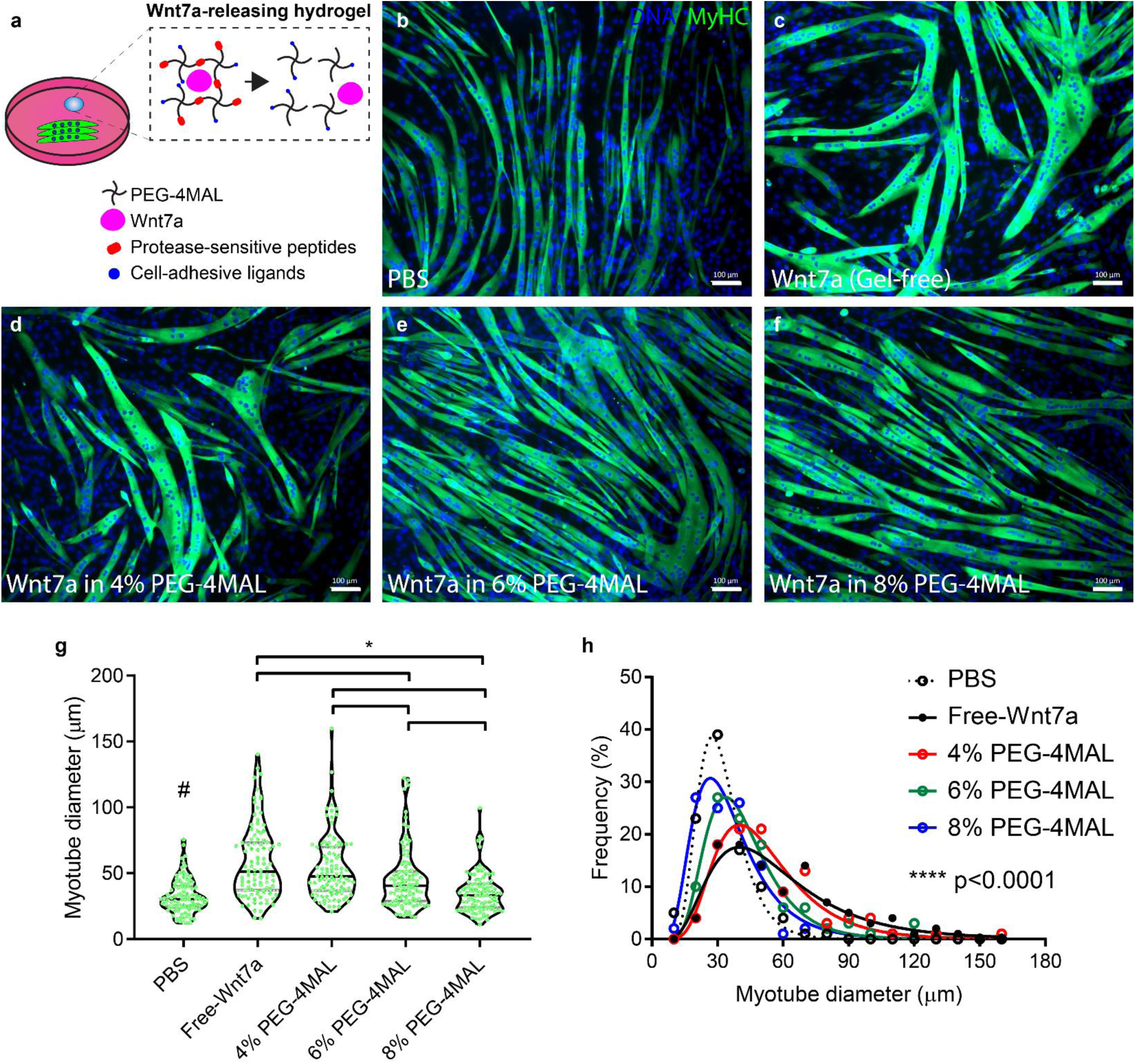
Hydrogel-released Wnt7a retains its bioactivity *in vitro*. **(a)** Schematic diagram of the experiment. Differentiating C2C12 myotubes treated with **(b)** PBS, **(c)** Wnt7a (gel-free), **(d)** Wnt7a in 4% PEG-4MAL hydrogel, **(e)** Wnt7a in 6% PEG-4MAL hydrogel, and **(f)** Wnt7a in 8% PEG-4MAL hydrogel. Day 5. Scale bar 100 μm. **(g)** Violin plot of myotube diameter. n=100 myotubes/condition. Kruskal-Wallis test with Dunn’s multiple comparisons test. #*p*<0.001 vs. free-Wnt7a, 4% PEG-4MAL, and 6% PEG-4MAL. \**p*<0.05. **(h)** Histogram of myotube diameter distribution with the log-Gaussian fit. Extra sum of squares F-test for comparing curve fit. \*\*\*\**p*<0.0001.

### Engineered PEG-4MAL hydrogel enables effective local delivery of Wnt7a to the injured skeletal muscle

The PEG-4MAL hydrogel can adhere to the tissue through maleimide-thiol interactions at physiologic conditions, and thus offers a unique advantage for locally administering therapeutics at the site of tissue injury where direct injection is not practical (Fig. 3a). Indeed, retention of unpolymerized, reacting PEG-4MAL macromers on the surface of TA muscle is significantly higher compared to unpolymerized macromers with quenched maleimides (*p*<0.05) and untreated (*p*<0.01; Fig. 3b and c), corroborating the feasibility of adhering a Wnt7a-loaded PEG-4MAL hydrogel to skeletal muscle.

**Figure 3.**
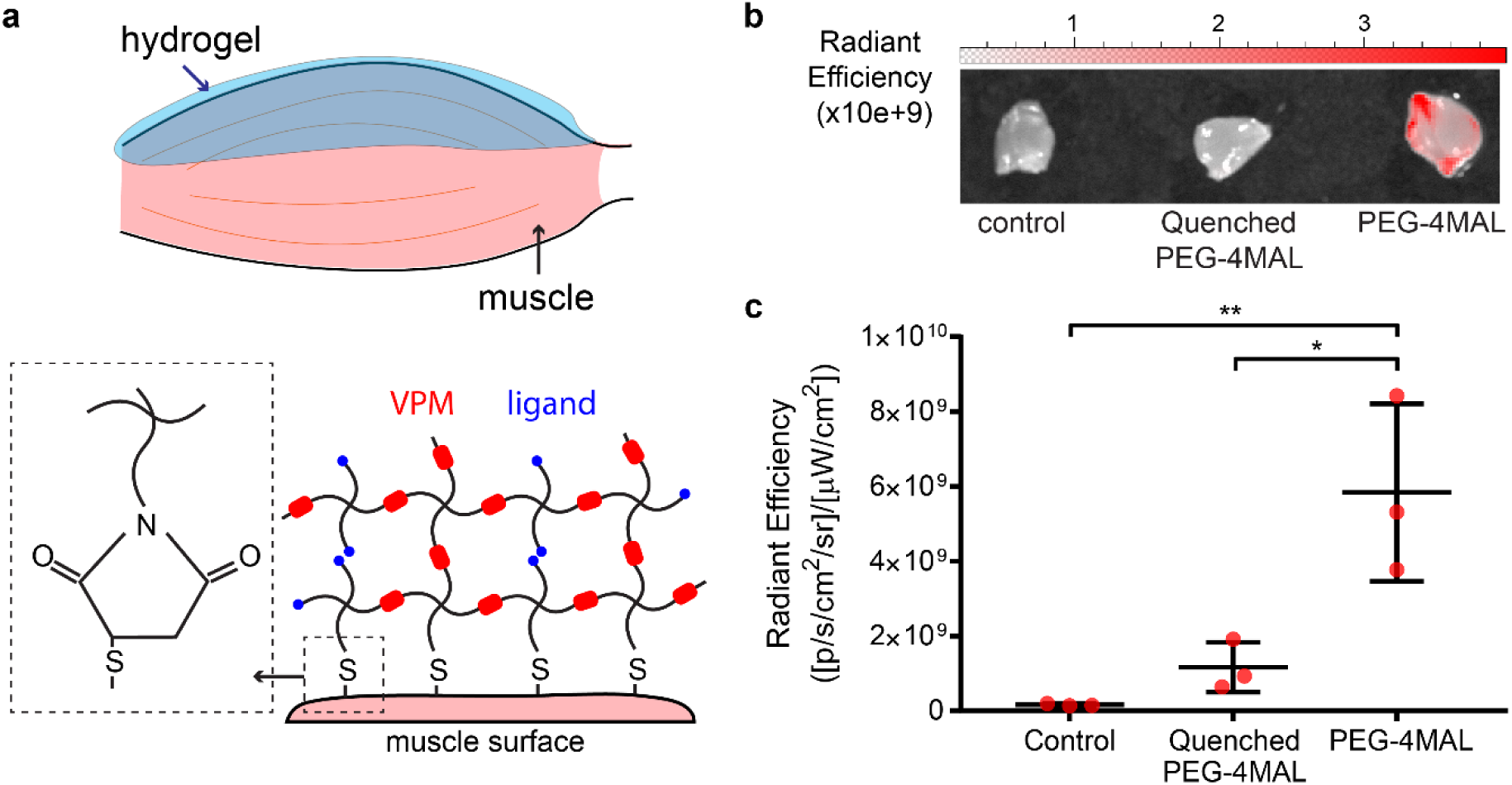
PEG-4MAL macromer adheres to the muscle surface. **(a)** Schematic diagram illustrating the ability of PEG-4MAL hydrogel to adhere to the muscle surface via maleimide-thiol reaction. **(b)** Representative IVIS images of muscles treated control (saline), quenched PEG-4MAL macromers conjugated with DyLight™-755-labeled RDG peptides, and PEG-4MAL macromers conjugated with DyLight™-755-labeled RDG peptides. **(c)** Quantification of radiant efficiency. One-way ANOVA with Tukey’s multiple comparisons test. \**p*<0.05, \*\**p*<0.01.

To further assess the feasibility of promoting muscle regeneration by locally administering Wnt7a in severe muscle injury, we delivered Wnt7a-loaded hydrogels to the supramuscular locus of freeze-injured TA muscle. The muscles treated with Wnt7a-loaded hydrogels exhibited significantly increased median myofiber area at the site of injury compared to the muscles treated with Wnt7a-free hydrogels (Fig. 4a and b; *p*<0.05). Treatment with Wnt7a-loaded hydrogels also significantly increased TA mass compared to TAs treated with Wnt7a-free control hydrogels (Fig. 4c; *p*<0.01). Furthermore, a significantly higher number of endogenous Pax7^+^ MuSCs were found in the TAs treated with Wnt7a-loaded hydrogels compared to control gels (Fig. 4d and e; *p*<0.05). Collectively, these results suggest that hydrogel-release Wnt7a retains its bioactivity *in vivo*, and that local delivery of Wnt7a to severely injured muscle using the engineering PEG-4MAL hydrogel is feasible and effective in accelerating muscle regeneration.

**Figure 4.**
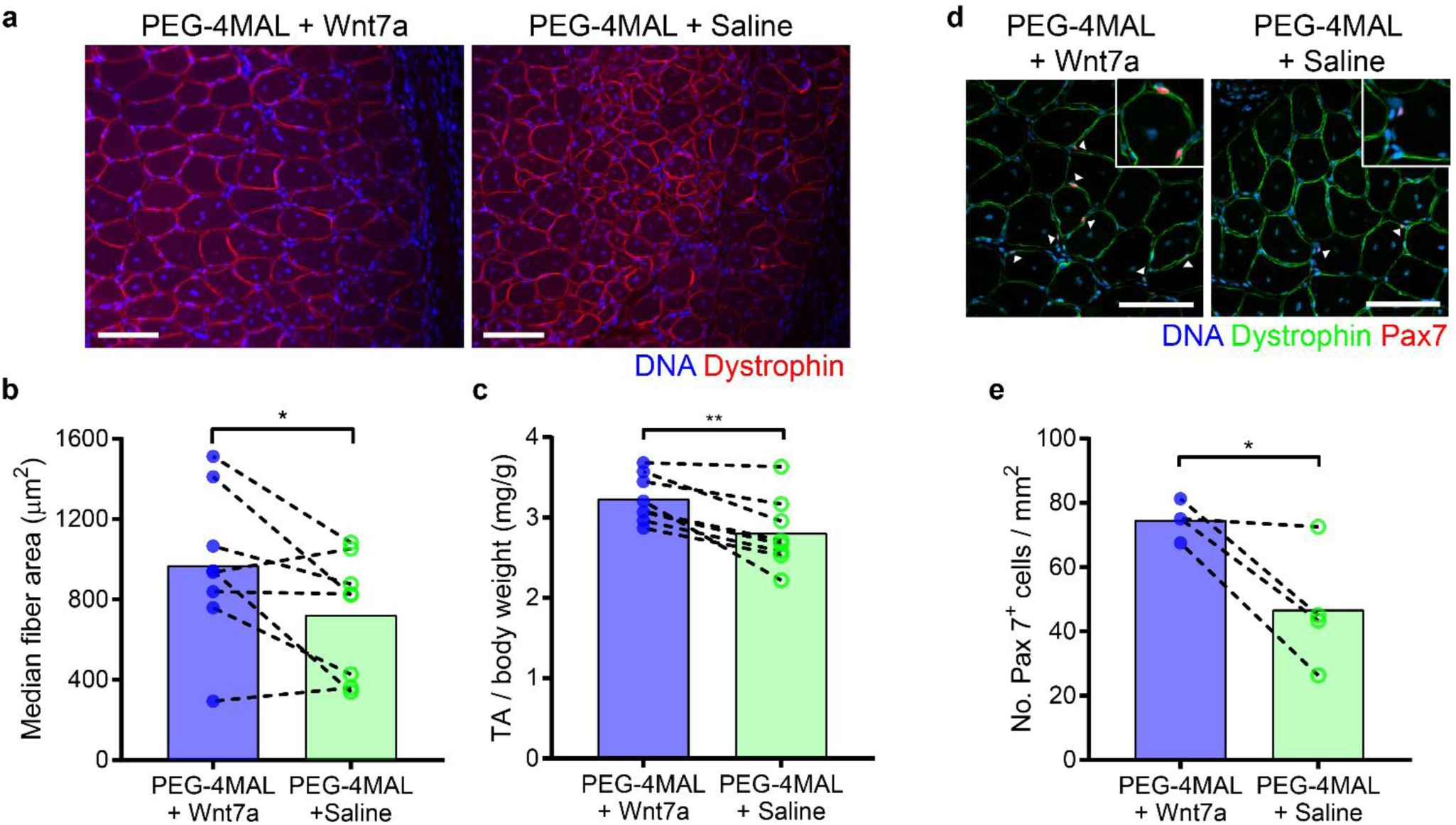
Engineered PEG-4MAL hydrogel enables effective local delivery of Wnt7a to the injured skeletal muscle. **(a)** Representative images of TA muscles treated with PEG-4MAL hydrogels loaded with Wnt7a or saline. Day 14 post-treatment. Scale bar 100 μm. **(b)** Median fiber area. Day 14 post-treatment. Two-tailed paired t-test. **(c)** Mass of TA muscles normalized to the respective body weight. Day 14 post-treatment. Two-tailed paired t-test. **(d)** Representative images of TA muscles treated with PEG-4MAL hydrogels loaded with Wnt7a or saline. Arrows indicate Pax7^+^ muscle stem cells. Day 14 post-treatment. Scale bar 100 μm. **(e)** The number of Pax7^+^ muscle stem cells per mm^2^. Day 14 post-treatment. Two-tailed paired t-test. \**p*<0.05, \*\**p*<0.01.

### Wnt7a promotes MuSC migration through the engineered PEG-4MAL hydrogel

Wnt7a promotes symmetric expansion and migration of MuSCs through the planar cell polarity pathway involving Dvl2 and the small GTPase Rac1 (6, 7, 10). Furthermore, brief treatment of MuSCs prior to transplantation enhances engraftment (7), supporting the application of hydrogel-mediated MuSCs transplantation with Wnt7a to treat muscle trauma. We have previously engineered a PEG-4MAL hydrogel that is conducive to primary MuSC function and demonstrated that MuSC transplantation using the engineered hydrogel significantly improves the engraftment potential (16). To determine how Wnt7a influences MuSC proliferation within the engineered hydrogel *in vitro*, we encapsulated freshly isolated MuSCs in hydrogels presenting either RGD or RDG (scrambled inactive control) integrin-targeting ligands with or without Wnt7a. Myogenic colonies formed in the RGD-presenting hydrogel were significantly larger than the colonies formed in the RDG-presenting hydrogels (Fig. 5a and b; *p*<0.001). However, encapsulated Wnt7a did not result in an increased size of myogenic colony formation in both RGD-and RDG-functionalized hydrogels (Fig. 5a and b). Although no significant difference in EdU incorporation was observed within the RGD-presenting gels with encapsulated Wnt7a, the number of EdU^+^ nuclei per colony in the Wnt7a-treated RGD-presenting hydrogel was significantly higher than both RDG-functionalized conditions (Fig. 5a and c; *p*<0.01). Furthermore, no difference in colony density was observed in RGD-presenting hydrogels with encapsulated Wnt7a compared to control RGD-presenting gels (Fig. 5d and e), indicating that Wnt7a does not increase the proliferation capacity of myogenic progenitors in the engineered PEG-4MAL hydrogel. However, MuSCs seeded on top of Wnt7a-releasing hydrogel exhibited increased invasion distance into the hydrogel compared to MuSCs seeded on Wnt7a-free hydrogel (Fig. 5f and g). This result shows that Wnt7a promotes MuSCs migration through the engineered synthetic matrix, which could further improve transplantation efficacy when delivered to the injured muscle *in vivo*.

**Figure 5.**
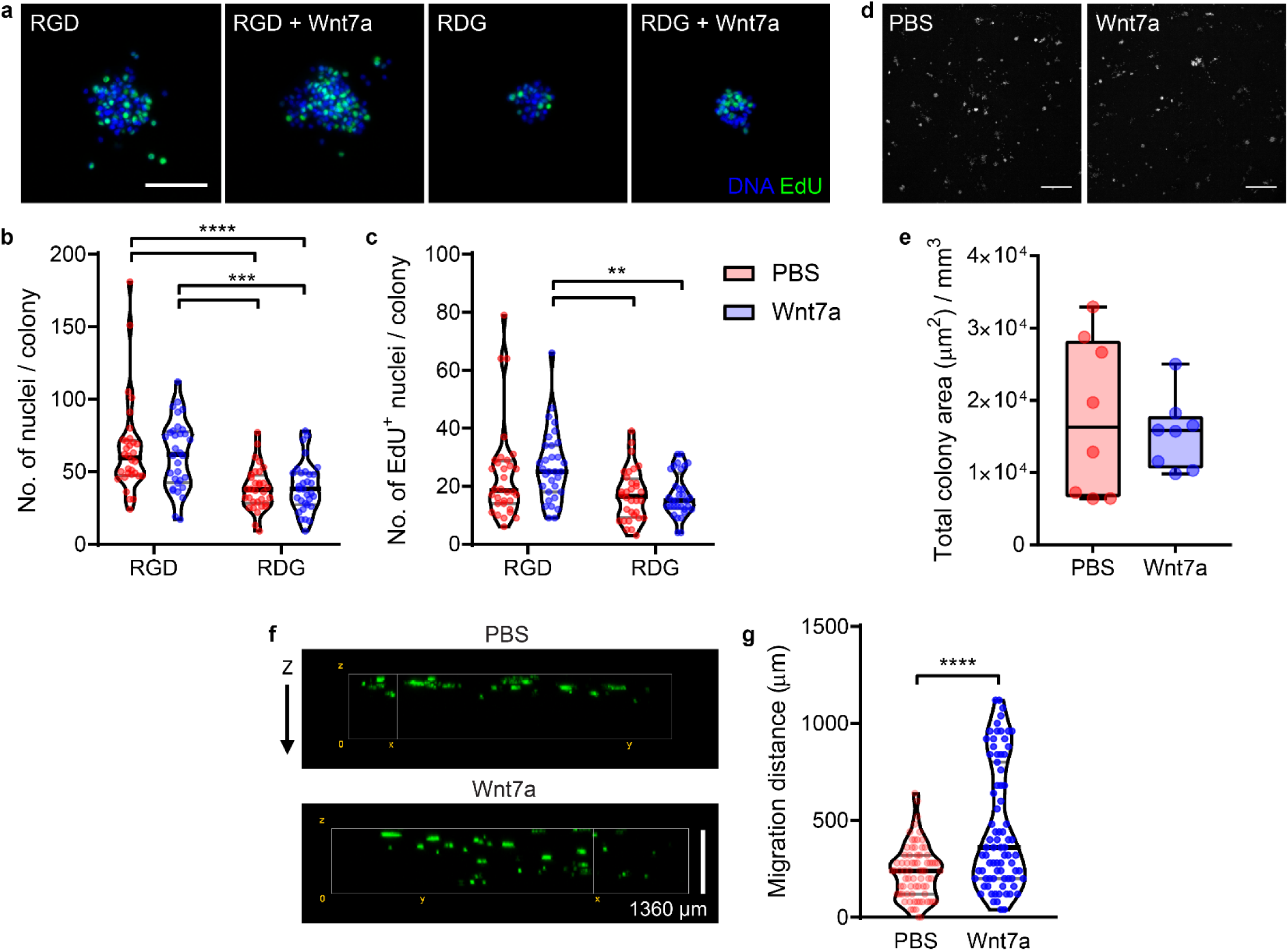
Wnt7a does not increase MuSC proliferation but promotes muscle stem cells migration through the engineered PEG-4MAL hydrogel. **(a)** Representative confocal images of myogenic colonies formed in RGD-presenting PEG-4MAL hydrogel, RGD-presenting PEG-4MAL hydrogel with Wnt7a, RDG-presenting PEG-4MAL hydrogel, and RDG-presenting PEG-4MAL hydrogel with Wnt7a. Scale bar 50 μm. **(b)** Violin plot of myogenic colony size. Two-way ANOVA with Sidak’s multiple comparisons test. n=32 colonies/condition. **(c)** Violin plot of EdU quantification. Two-way ANOVA with Sidak’s multiple comparisons test. n=32 colonies/condition. **(d)** Representative confocal images of nuclei-stained myogenic colonies formed in RGD-presenting PEG-4MAL hydrogels with or without Wnt7a. Scale bar 500 μm. **(e)** Colony density quantified by total colony area per volume. **(f)** 3D projection of GFP+ muscle stem cells migrating through the hydrogel from the top surface. **(g)** Violin plot of cellular migration distance. Day 3. n=79-74 cells/condition. Mann-Whitney test. \*\**p*<0.01, \*\*\**p*<0.001, \*\*\*\**p*<0.0001.

### Wnt7a promotes MuSC migration *in vivo*

Delivery of MuSCs using the engineered PEG-4MAL hydrogel to the supramuscular locus of freeze-injured TA muscle *(mdx)* significantly increases engraftment potential compared to the hydrogel-free condition (Fig. 6a and b), consistent with our previous findings (16). To further evaluate the effect of Wnt7a on MuSC engraftment, we next delivered freshly isolated GFP^+^ MuSCs using the engineered PEG-4MAL hydrogels with or without encapsulated Wnt7a to the supramuscular locus of freeze-injured TA muscles. Transplantation of MuSCs via the engineered PEG-4MAL hydrogel containing Wnt7a did not increase the number of GFP^+^ fibers engrafted (Fig. 6c and d; *p*>0.05). However, the minimal distance of GFP^+^ engrafted fibers from the TA surface was significantly increased when MuSCs were co-delivered with Wnt7a via the PEG-4MAL hydrogel (Fig. 6c and e), suggesting that MuSCs delivered in the absence of Wnt7a preferentially engraft along the supramuscular locus, while MuSCs co-delivered with Wnt7a engraft at a deeper location from the TA surface and the delivery site. Taken together, the data demonstrate that co-delivery of MuSCs and Wnt7a to the supramuscular locus of injured TA muscle using the engineered PEG-4MAL hydrogel promotes MuSC *in vivo* migration without impacting the engraftment efficiency.

**Figure 6.**
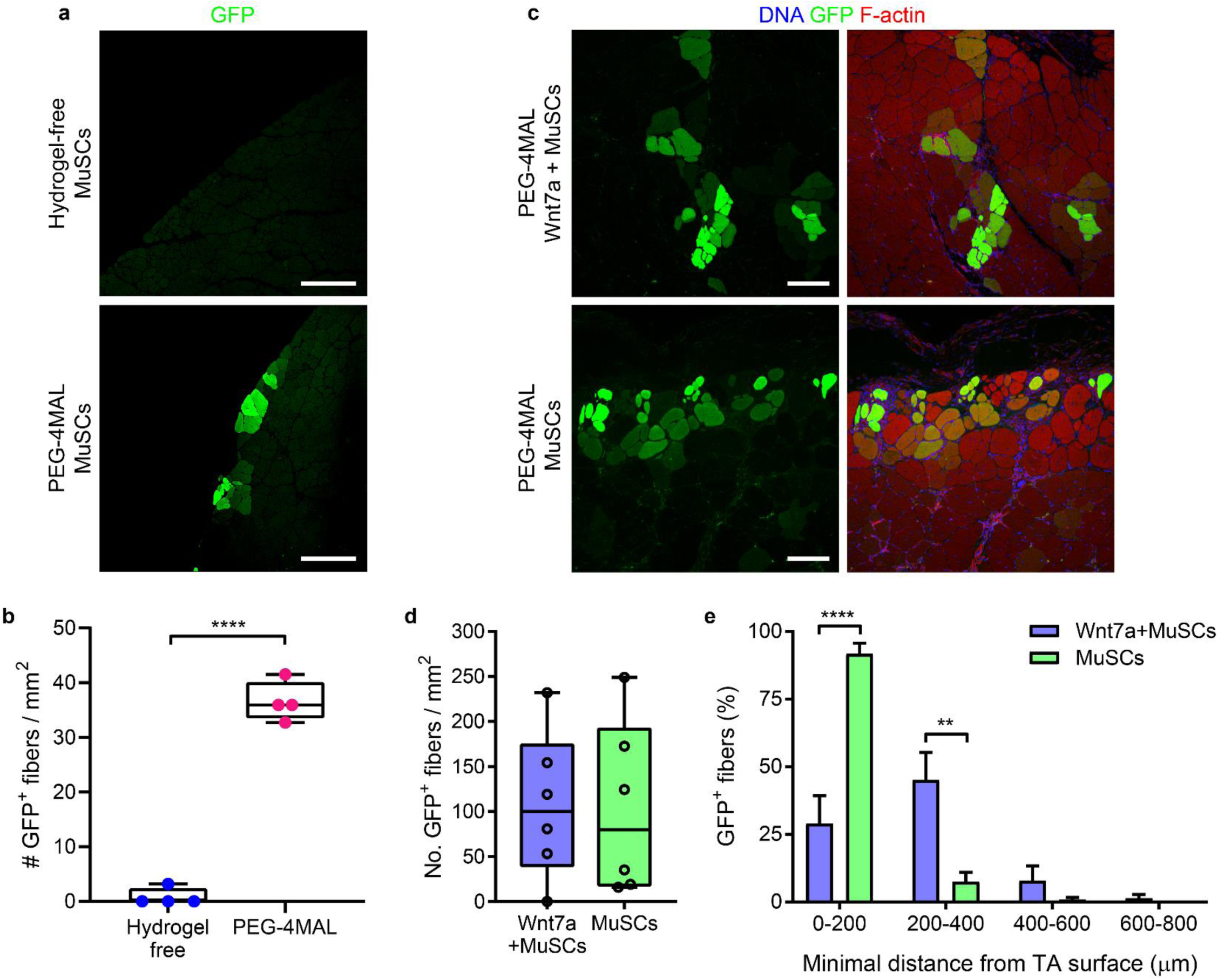
Wnt7a increases migration potential of PEG-4MAL-released muscle stem cells *in vivo*. **(a)** Representative confocal images of TA muscles treated with muscle stem cells alone and muscle stem cells delivered using the engineered PEG-4MAL hydrogel. 40,000 cells delivered per TA. Scale bar 250 μm. **(b)** The number of engrafted GFP^+^ fibers. Two-tailed unpaired t-test. **(c)** Representative confocal images of TA muscles treated with muscle stem cells encapsulated in PEG-4MAL hydrogels with or without Wnt7a. 50,000 cells delivered per TA. Scale bar 100 μm. **(d)** The number of engrafted GFP^+^ fibers. Two-tailed paired t-test. **(e)** Histogram of GFP^+^ fibers (%) as a function of minimal distance from the TA surface. Two-way repeated measures ANOVA with Sidak’s multiple comparisons test. \*\**p*<0.01, \*\*\*\**p*<0.0001.

## DISCUSSION

MuSC transplantation and pro-myogenic factor delivery are effective therapeutic strategies to regenerate damaged and diseased skeletal muscles, but direct delivery of cells and proteins to severely injured (e.g., laceration and volumetric muscle loss) muscles is challenging due to a loss in tissue integrity. To address this challenge, we engineered a synthetic PEG-4MAL hydrogel to facilitate the co-delivery of MuSCs and Wnt7a protein in conditions where direct injection may not be feasible. The maleimide groups functionalized on the PEG macromer efficiently react with thiol-presenting biological tissues to form stable covalent bonds via Michael-type addition, enabling hydrogel adhesion to the site of tissue injury and subsequently permit local delivery of cells and therapeutics (Fig. 3). We have previously demonstrated that the engineered PEG-4MAL hydrogel supports primary MuSC survival, proliferation, and differentiation, and enables delivery of MuSCs in dystrophic and aged skeletal muscles with comorbid trauma (16). The engineered PEG-4MAL hydrogel significantly improves the cellular engraftment efficiency compared to hydrogel-free and collagen gel controls (16). In this study, we further demonstrate that the engineered PEG-4MAL hydrogel enables controlled delivery of Wnt7a to accelerate the regenerative response by targeting the endogenous MuSCs and myofibers (Fig. 4). In addition, we establish the feasibility of co-delivering exogenous MuSCs and Wnt7a using the engineered PEG-4MAL hydrogel (Fig. 6). Co-delivery of MuSCs and Wnt7a significantly increases the migration capacity of the donor MuSCs *in vivo* (Fig. 6), suggesting that the function of hydrogel-released MuSCs can be modulated by incorporation of pro-myogenic proteins and growth factors.

Hydrogel-mediated delivery of cells and protein-based therapeutics offer numerous advantages over vehicle-free cell and protein delivery methods (13, 15). In protein and drug delivery, hydrogels enable spatiotemporally controlled release of therapeutics and minimize the need for the repeated administration of high dosages. Hydrogel-based delivery also permits localized administration of therapeutics to the target tissue of interest, reducing the risk of systemic off-target effects. In the current study, we show that the release rate of Wnt7a from the engineered PEG-4MAL hydrogel can be tuned by modulating the macromer density and protease-mediated degradation of the hydrogel (Fig. 1a). Delivery of Wnt7a-loaded PEG-4MAL hydrogel to the injured muscle significantly accelerated the regenerative response at the injury site by day 14 (Fig. 4). Although beyond the scope of the current study, different muscle types (e.g., craniofacial, diaphragm), injuries (e.g., volumetric muscle loss, cardiotoxin), and diseases (e.g., Duchenne muscular dystrophy) may require context-specific release kinetics of Wnt7a to achieve desired therapeutic effects. Furthermore, using the PEG-4MAL hydrogel platform, physical and biochemical properties, including matrix stiffness, cell-adhesive ligand presentation, structure, and degradability, can be systematically modulated to create a cell-instructive microenvironment (16, 25). Here, we demonstrate that the engineered PEG-4MAL hydrogel supports the proliferation, and migration of primary MuSCs in the presence of Wnt7a (Fig. 5). Local co-delivery of MuSCs and Wnt7a using the engineered PEG-4MAL hydrogel to the injured muscle is also feasible (Fig. 6). In subsequent studies, it would be important to evaluate the efficacy of this strategy in other clinically-relevant models of muscle injuries and diseases.

Wnt7a induces myofiber hypertrophy through the Akt/mTOR pathway (8) and symmetric expansion of MuSCs through the planar cell polarity pathway (6, 10). Furthermore, pre-treatment of MuSCs with Wnt7a stimulates cell migration and engraftment potential, ultimately leading to significantly increased muscle function (7). In this study, we show that Wnt7a released from the hydrogel retains its bioactivity, promotes myofiber hypertrophy, and increases the number of Pax7^+^ MuSCs present at the injury site (Fig. 2 and 4). Co-delivery of MuSCs with Wnt7a using the engineered hydrogel also increased the migration distance from the delivery site, suggesting that Wnt7a also increases the MuSC dispersion during the engraftment process when delivered using the hydrogel (Fig. 6c and e). In addition, although previous work has demonstrated that Wnt7a treatment significantly increases the engraftment potential of MuSCs (7), co-delivery of MuSCs with Wnt7a had an insignificant effect on the number of GFP^+^ fibers engrafted (Fig. 6c and d). This suggests that the differences in the injury model (*e.g.*, cardiotoxin *vs.* freeze injury), mode of delivery (e.g., direct injection *vs.* supra-muscular casting), and use of cell delivery vehicle may influence how the cells engraft into the host muscle upon transplantation.

In summary, we engineered a synthetic PEG-4MAL hydrogel that facilitates the co-delivery of Wnt7a and MuSCs for accelerated skeletal muscle healing in conditions where direct intramuscular injection may be challenging. The maleimide-thiol interaction enables adhesion of Wnt7a and MuSC-encapsulated hydrogel to the freeze-injured muscle surface, and the hydrogel-encapsulated Wnt7a significantly increases muscle fiber hypertrophy, endogenous MuSC expansion, and exogenous MuSC migration during the engraftment process. This work establishes a proof-of-concept biomaterial-mediated therapeutic strategy for promoting skeletal muscle regeneration in the context of traumatic muscle injuries and neuromuscular diseases.

## ACKNOWLEDGMENTS

We thank the Physiological Research Laboratory and core facilities at the Parker H. Petit Institute for Bioengineering and Bioscience at the Georgia Institute of Technology for the use of shared equipment and services. Research reported in this publication was supported by the National Institutes of Health under Award Numbers F32HL140821, R01AR062368, and R21AR072287. The content is solely the responsibility of the authors and does not necessarily represent the official views of the National Institutes of Health. The authors declare no competing interests.

## AUTHOR CONTRIBUTIONS

W. M. Han, A. J. García, and Y. C. Jang designed the study. W. M. Han, M. Mohiuddin, and S. E. Anderson performed the experiments. W. M. Han, A. J. García, and Y. C. Jang wrote the manuscript. All authors reviewed the manuscript.

